# Increased collective migration correlates with germline stem cell competition in a basal chordate

**DOI:** 10.1101/2023.07.11.548629

**Authors:** Megan K. Fentress, Anthony W. De Tomaso

## Abstract

Cell competition is a process that compares the relative fitness of progenitor cells, resulting in ‘winners’, which contribute further to development, and ‘losers’, which are excluded, and is likely a universal quality control process that contributes to the fitness of an individual. Cell competition also has pathological consequences, and can create super-competitor cells responsible for tumor progression. We are studying cell competition during germline regeneration in the colonial ascidian, *Botryllus schlosseri*. Germline regeneration is due to the presence of germline stem cells (GSCs) which have a unique property: a competitive phenotype. When GSCs from one individual are transplanted into another, the donor and recipient cells compete for germline development. Often the donor GSCs win, and completely replace the gametes of the recipient-a process called germ cell parasitism (gcp). gcp is a heritable trait, and winner and loser genotypes can be found in nature and reared in the lab. However, the molecular and cellular mechanisms underlying gcp are unknown. Using an ex vivo migration assay, we show that GSCs isolated from winner genotypes migrate faster and in larger clusters than losers, and that cluster size correlates with expression of the Notch ligand, Jagged. Both cluster size and jagged expression can be manipulated simultaneously in a genotype dependent manner: treatment of loser GSCs with hepatocyte growth factor increases both jagged expression and cluster size, while inhibitors of the MAPK pathway decrease jagged expression and cluster size in winner GSCs. Live imaging in individuals transplanted with labeled winner and loser GSCs reveal that they migrate to the niche, some as small clusters, with the winners having a slight advantage in niche occupancy. Together, this suggests that the basis of GSC competition resides in a combination in homing ability and niche occupancy, and may be controlled by differential utilization of the Notch pathway.

## Introduction

Cell competition is a process that compares the relative fitness of progenitor cells, and results in healthier cells contributing a higher proportion to the final tissue composition, while damaged, stressed or otherwise suboptimal cells are removed. First described in Drosophila in the 1970s [1], cell competition has been demonstrated in organisms that range from the social amoebae Dictyostelium, to humans, and is likely a universal quality control mechanism that plays a role both during development, and tissue homeostasis in adults (reviewed in [2]). Cell competition is dependent on cells competing for extrinsic survival factors [3], or comparing some intrinsic limiting resource, such as rates of translation, growth, or expression levels of certain proteins (e.g., Myc, p53), to neighboring cells and responding to the status of their neighbors. The outcome of competition varies with the organism and tissue studied. For example, in the developing Drosophila wing, winner cells instruct loser cells to commit apoptosis [2]. In contrast, studies focused on Drosophila ovarian stem cells have shown that competition occurs for niche occupancy via differences in cell adhesion. The winner cell displaces the loser cell from the germ cell niche, and concomitant instructive cues for self-renewal and/or differentiation-thus losers are excluded from contribution to further development [4].

These kinds of competitive events have also been observed in adult stem cell populations in mammals. For example, it has been shown that successful engraftment of transplanted hematopoietic stem cells involves competition between the donor and recipient cells for niche occupancy, with successful engraftment correlating to levels of p53 expression [5]. In addition, competition is likely involved in pathological processes such as cancer. When genes involved in competition are overexpressed, they can create super-competitors that overgrow normal cells (reviewed in [6]), thus competitive interactions likely underlie tumor cell selection and progression. In summary, cell competition is a phenomenon observed from single cell organisms to humans, and plays a role in multiple processes.

Despite its ubiquity, the molecular mechanisms underlying cell competition are not well understood. For example, differences in expression of a number of genes correlate to the competitive interactions between cells, including myc, p53, and wnt; but it is not known if there is a common downstream regulatory mechanism that interpret those measurements [2,7]. In addition, competition is context dependent; cells are comparing themselves to nearby cells, thus even the concept of cell fitness is difficult to define, and not yet quantifiable. Most importantly, the examples above are interactions between somatic cells, but it is an individual that is under selection, not just the competitive interactions. As such, there could be multiple redundant mechanisms contributing to competition that will be difficult to separate out without genetic variants to compare.

We are studying cell competition in a novel model organism, the colonial ascidian, *Botryllus schlosseri*. Ascidians are marine organisms which grow in shallow waters throughout the world, and are part of the Tunicata, considered the basal chordates [8]. Embryogenesis results in a chordate tadpole larva that commences a short swimming phase, culminating in the identification of a suitable substrate for the adult form. The larva then settles and undergoes metamorphosis into a sessile and transparent adult body plan, called an oozooid. The oozooid is a filter-feeding individual with a complex anatomy, including a GI tract, heart, both a central and peripheral nervous system, a complex musculature and hematopoietic system. Botryllus belongs to a subset of ascidians that have a colonial life history-meaning that an individual does not grow by increasing in size, but rather by a lifelong, recurring asexual budding process during which entire bodies are regenerated de novo every week. This results in a constantly expanding colony of genetically identical individuals, called zooids, with the same body plan as the oozooid. All zooids are connected by a common extracorporeal vasculature, which ramifies throughout the colony. At the periphery of the colony, these blood vessels terminate in blind ended protrusions called ampullae (Supplemental Figure 1). While all zooids are connected by a common vasculature, they are not dependent on each other. In the lab, colonies grow on glass slides, and portions of a colony can be surgically removed, transferred to a new slide, and will continue to grow-these are called subclones, and this can be done multiple times. In summary, independent experiments can be carried out using the exact same genotype.

Each week, each zooid initiates the regeneration of 1-4 new buds, during which all somatic and germline tissues develop de novo in a synchronized fashion throughout the colony. The best understood regenerative process in Botryllus occurs in the germline. Like most metazoans, Botryllus sets aside a population of PGCs early in embryogenesis [9]. However, unlike most model organisms, Botryllus retains a population of mobile, self-renewing, lineage-restricted adult germline stem cells (GSCs) that retain pluripotency for life of the individual, which ranges from 6 months to > 2 years in the lab [9,10]. Every week a subset of these GSCs settle and differentiate into gametes, while others self-renew, and will migrate to the newly forming niches in the subsequent generation of developing bodies. Migration occurs over a defined 48 hr. period, and during this window GSCs are also found in the colony vasculature and ampullae [11]. So, while in most model organisms PGC migration occurs once during embryogenesis, in Botryllus a population of long-lived, lineage restricted, self-renewing GSCs migrate synchronously from an old niche to a new niche during a defined 48-hour period each and every week during the life of an individual (Supplemental Figure 1; [11]).

As individuals grow by asexual reproduction, they continuously expand over the substrate, and when two grow into contact, the first tissues to touch are the ampullae. This contact initiates a natural transplantation reaction that can result in either fusion of the two ampullae, forming a parabiosis between the two individuals- or an inflammatory rejection response that blocks parabiosis (Supplemental Figure 1). Fusion or rejection is controlled by a single highly polymorphic locus, called the *fuhc*, and two individuals fuse if they share one or both fuhc alleles, and reject if no alleles are shared between them [12]. Following fusion of two compatible individuals, the GSCs can migrate from one genotype to the other via the parabiotic linkage. When GSCs from two individuals co-exist in a single body, they will compete for germline development. In some cases, GCSs from one genotype will win, and only that genotype will contribute to the germline of all of the zooids from both individuals- and this dominance will remain for years following transplantation, even if the two colonies are surgically separated (Supplemental Figure 1 [13–16]. This competitive trait, called germ cell parasitism (gcp), is both repeatable in independent pairings of the same genotypes, and heritable. There are winner and loser genotypes found in both natural populations and reared in our lab [10,14].

If fusion occurs between two genotypes with equal competitive abilities, GSCs from both genotypes will contribute to the mature gametes throughout the chimera (Supplemental Figure 2). Both GSCs and blood progenitors are mobile and can migrate between two individuals in a parabiotic pair, but the rest of the somatic tissues cannot [17,18]. Thus, a zooid can consist of somatic tissues from one genotype, but be making the gametes of the other.

Importantly, the parasitic phenotype is autonomous to the GSCs, as their competitive phenotype is retained during experimental transplantation [10]. GSCs can be isolated by FACS and transplanted into recipient individuals and their contribution to gamete development monitored. If GSCs are isolated from a winner genotype, and transplanted into a loser, they contribute to germline development in the recipient. In contrast, if GSCs from a loser genotype are transplanted into a winner, they do not [10]. Using serial transplantation, we have also found that GSCs are lineage restricted and self-renewing, but how these long-lived progenitors compete for germline development is not well understood [10]. Here we have utilized the natural genetic variability in gcp to study the cellular and molecular mechanisms which underlie stem cell competition.

## Materials and Methods

### Animal husbandry identification of winner/loser pairs

*B. schlosseri* colonies used in this study were lab-cultivated strains, spawned from individuals collected in Santa Barbara, CA, cultured in laboratory conditions at 18–20 °C, and staged as described in detail previously [19]. Winner/loser pairs are defined as 1) genotypes which are histocompatible and naturally fuse to form parabiotic pairs; and 2) show complete germline parasitism, i.e., following fusion all mature gonads are descendant from only one genotype. In addition, parasitism must be repeatable (consistent among multiple pairings of naïve subclones of each genotype) and stable [10,14]. Here we assayed chimerism of testes, as they can be easily isolated with no somatic tissue contamination. To identify winner/loser pairs, we used large healthy genotypes that could be repeatably separated into pieces (also called subcloning, the independent pieces are called subclones). Genotypes were visually tested for fusion by placing subclones in direct contact on glass slides. If colonies were compatible and fused, the parabiotic chimeras were maintained in a separate tank and allowed to go through at least 6 blastogenic cycles, then individual testis (n =10) were genotyped. Three independent experiments were done for each fusible genotype pair, and 3 winner/loser pairs of individuals were identified (i.e., all testis were from one genotype) and used in this study. Parasitism was observed in 15% of the fusible pairs tested here. To normalize for any potential genotype specific results, in each experiment in this study, each pair was used interchangeably at least once, and the data was aggregated. No significant differences were found.

### Genotyping

For genotyping, we used codominant length and dominant presence/absence polymorphisms in 10 polymorphic loci (Supplemental Figure 2). Initially, somatic tissue from naïve subclones of each genotype were characterized for polymorphisms for each locus, and specific combinations of loci were identified which could uniquely identify each genotype used in this study. Parabiotic pairs were tested at 6 and 12 weeks following fusion. A subclone was removed from the chimera was placed in a shallow dish of 70% ethanol. Zooids were sliced open and individual testis isolated under a dissecting microscope. Somatic tissue and blood were removed using small tweezers and repeated dipping into the ethanol until visibly clean. Genomic DNA was extracted using the Nucleospin DNA XS kit (Machery Nagel) following the manufactures instructions. Primers are listed below, and PCR conditions were 5 min at 95° C, followed by 36 cycles of 95° for 20 seconds, 56° C for 20 seconds and 72° C for 45 seconds. Samples were separated by gel electrophoresis using a 3% MetaPhor agarose (Lonza) using standard techniques. Individual testes contain about 10^5^ sperm, and as can be seen in previous studies [10,15–17] and Supplemental Figure 2, individual testis are clonally derived.

### Cell Sorting

FACS-based enrichment of GSCs were performed as previously described [10,20]. Briefly, genetically identical, stage-matched animals were pooled, and a single-cell suspension was generated by mechanical dissociation. Whole animals were minced and passed sequentially through 70μm and 40μm cell strainers in ice-cold sorting buffer (filtered sea-water with 2% horse serum and 50mM EDTA). FACS was performed using a FACSAria (BD Biosciences) cell sorter. Samples were gated with two previously identified markers for Botryllus germ cells, ALDH activity (detected using Aldefluor (Stem Cell Technologies)) and a monoclonal antibody to integrin alpha 6 (Anti-Human/Mouse-CD49f-dFlour450 (Ebioscience, San Diego, CA USA, clone G0H3)). Analysis was performed using FACSDiva software (BD Biosciences). Cells were sorted using a 70μm nozzle and collected into dissociation buffer.

### Quantitative RT–PCR

Sorted cells were pelleted at 700Xg for 10 min, and RNA was extracted using the Nucleospin RNA XS kit (Macherey Nagel), which included a DNAse treatment step. RNA was reverse transcribed into cDNA using random primers (Life Technologies) and Superscript II Reverse Transcriptase (Life Technologies). Quantitative RT-PCR was performed using a LightCycler 480 II (Roche) and LightCycler DNA Master SYBR Green I detection (Roche) according to the manufacturer’s instructions. The thermocycling profile was 5 min at 95 °C, followed by 45 cycles of 95 °C for 10 s, 60 °C for 10 s. The specificity of each primer pair was determined by BLAST analysis (to the *Botryllus* transcriptome database [46]), as well as by melting curve analysis and gel electrophoresis of the PCR product. To control for amplification of genomic DNA, ‘no RT’-controls were used. Primer pairs were analyzed for amplification efficiency using calibration dilution curves. All genes included in the analysis had cycle threshold (CT) values <35. Relative gene expression analysis was performed using the 2^-ΔΔCT^ Method. The CT of the target gene was normalized to the CT of the reference gene *elongation factor 1 alpha (EF1α)*: ΔC_T_=C_T(target)_–C_T (EF1α)_. To calculate the normalized expression ratio, the ΔCT of the test sample (IA6-positive cells) was first normalized to the ΔCT of the calibrator sample (IA6-negative cells): ΔΔC_T_=ΔC_T(IA6-positive)_-ΔC_T(IA6-negative)_. Second, the expression ratio was calculated: 2-^ΔΔCT^=Normalized expression ratio. The result obtained is the fold increase (or decrease) of the target gene in the test samples relative to IA6-negative cells. Each qPCR was performed at least three times on cells from independent sorting experiments, and each gene was analyzed in duplicate in each run. The ΔCT between the target gene and EF1alpha was first calculated for each replicate and then averaged across replicates. The average ΔCT for each target gene was then used to calculate the ΔΔCT as described above. Data are expressed as averages of the normalized expression ratio (fold change). Standard deviations were calculated for each average normalized expression ratio. Statistical analysis was performed using a paired, two-sided Student’s t-test. **P<0.05.

### Cell Tracking

GSCs were isolated as above from winner and −1 animals and labeled with either CM-Dil (Loser) (Cell Tracker™, Thermo Fisher Scientific) or Syto59 (Winner) (Cyto™ Red Fluorescent Nucleic Acid Stain, Thermo Fisher Scientific.). Isolated cells were spun at 1500rpm for 5 minutes, washed in filtered sea water, and incubated in label for 2 hrs. They were the spun at 1500rpm for 5 minutes and resuspended at 20,000 cells/uL. Loser genotype colonies were microinjected with the labeled cells into the blood stream, 24 hours later colonies were imaged using an Olympus FLV1000S Spectral Laser Scanning Confocal at 20X.

### Transwell migration assays

Migration assays were done as published previously [20,21]. Briefly, transwell filters with 8μm pore size inserted in a 24-well plate (Corning) were coated with laminin over night at 4 °C and briefly air dried before adding 50,000 sorted cells, resuspended in 100 μl filtered sea water with 10% DMEM, 1% FBS. The bottom of the well contained filtered sea water with 10% DMEM/1% FBS, and vehicle (control) or S1P (2μM or 0.2μM). After 4hr incubation at room temperature, nuclei were stained with Hoechst 33342 and counted in images taken at three random locations at x100 magnification. Average cell coverage was analyzed using “cell count” in FIJI software. All assays were performed in triplicates with cells from three independent sorts, and each winner/loser pair was used in each assay at least once. Data from all genotypes were combined and statistical analysis was performed using a paired, two-sided Student’s t-test.

### Small-molecule inhibitor treatment

Isolated cells were incubated for 2 hours in 200 μl of filtered sea water with 10% DMEM, 1% FBS containing one of the following reagents: 10μM U0126, 10μM Ly294002 (Cell Signaling), 40ng/ml Hepatocyte Growth Factor (Sigma Aldrich) or 5μM FR180204 (Tocris). Controls were incubated in sea water without inhibitors plus vehicle (0.1% ethanol or 0.001% dimethylsulphoxide). Inhibitor doses were determined empirically and represent the lowest concentration that gave maximum results in dose-response curves. For each treatment, three genetically identical subclones of each winner/loser pair were treated simultaneously. Transwell Migration Assays and Quantitative RT-PCR was performed as described above and data are reported as averages from all experiments. Statistical analysis was performed using a paired, two-sided Student’s t-test (***p<0.001).

### Primers used in this study

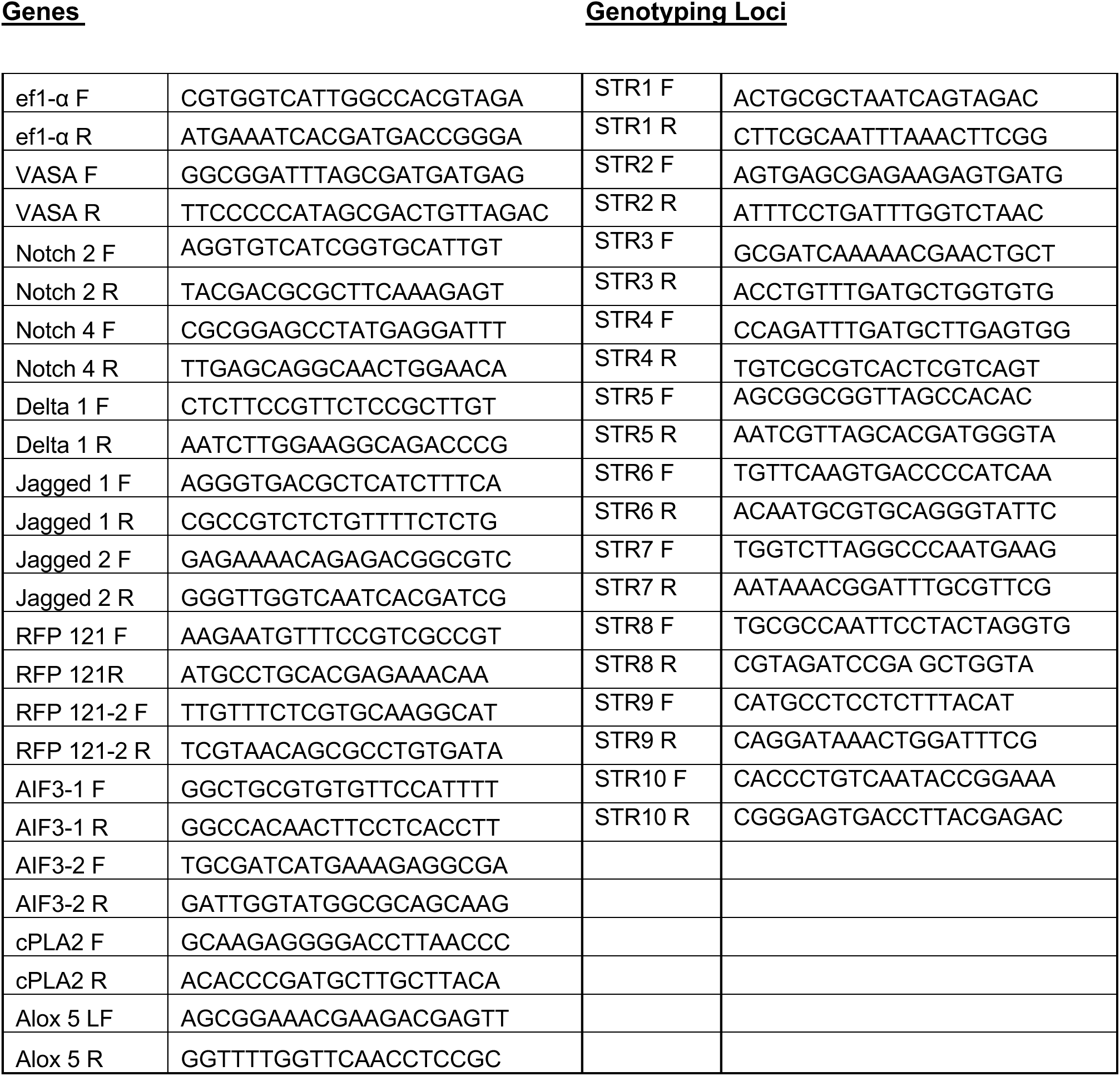

## Results

### Winner GSCs have increased migration capabilities compared to loser GSCs

Two broad and potentially interrelated hypothesises could explain the basis of germline parasitism: first, the winner GSCs could be faster at detecting, migrating to and occupying the developing germline niche; and/or there may be competition for maintaining access to the niche during gametogenesis. Using a combination of in vitro transwell migration assays and in vivo analyses, we had previously found that the migration of GSCs is controlled by a gradient of sphingosine-1-phosphate (S1P) secreted by the newly developing niche [20,21], and also that S1P chemotaxis may also be dependent on the generation of an autonomous gradient of the bioactive lipid, 12-S-HETE [21]. In order to characterize general migration mechanisms, these studies were done using multiple wild-type genotypes, without regard to the competitive phenotype. Here, we wanted to test whether winner and loser GSCs respond differently to S1P. We identified three genetically distinct pairs of winner/loser genotypes in our lab-reared cultures, isolated GSCs by FACS, and compared their migration to S1P gradients in vitro (described in the Methods section).

As shown in Figure 1, in all conditions, the number of winner GSCs migrating during the incubation period were higher than the loser GSCs-in other words, winner GSCs migrated faster than loser GSCs. Both genotypes revealed the equivalent dose-dependency to S1P previously observed [20], with an optimal migration to an S1P concentration of 2μM compared to a concentration of either 0.2 μM or 20 μM. Interestingly, winner GSCs also displayed slightly higher migration in unstimulated control experiments. In summary, the results from this experiment indicate that S1P induces chemotaxis in both winner and loser GSCs, however, winner-GSCs possess seem to possess an intrinsically faster migratory activity compared to loser cells (Figure 1).

**Figure 1:**
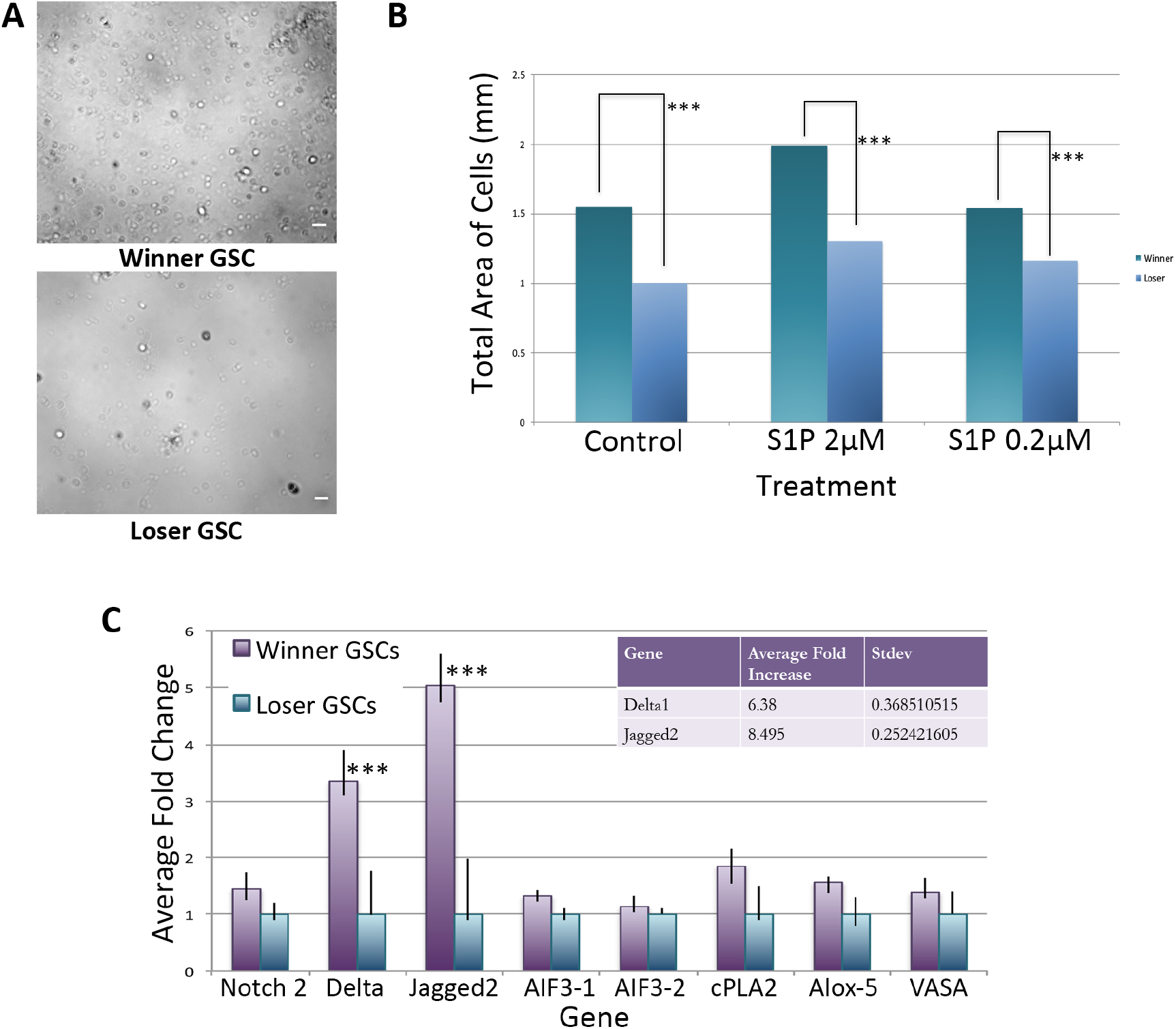
Winner GSCs have increased migration capabilities, and expression of jagged and delta compared to loser GSCs. **A.** Images of winner (top) and loser (bottom) GSCs after 4 h of migration in control treatment SB= 20 um. **B.** Plot of total area of cells following 4 h of migration. Winner GSCs (purple), loser GSCs (blue), treatments on X-axis. **C.** qRT-PCR analysis of isolated GSCs from winner and loser genotypes. Data expressed as averages of relative fold change normalized to loser GSCs. Each winner loser pair was assessed 3 times, and standard deviation calculated (n =9). ***P<0.001 using Students t-test.

As described in the Methods, we assess migration in this assay via nuclear staining of cells in the bottom of the transwell. Unexpectedly, in these experiments we found that following migration through the transwell filter, the majority of winner GSCs were found in larger clusters, while the loser GSCs migrated mostly as small clusters and some single cells (Figure 5 A,D). Interestingly, the presence of S1P slightly increased cluster size in both winner and loser GSCs vs unstimulated controls (Figure 5), and these observations were consistent between all three winner/loser genotypes. In summary, GSCs, which are FACS sorted as single ALDH+/IA6+ cells, appear to coalesce into clusters during incubation and migration, with winners forming larger clusters than losers. In addition, cluster size correlates to the speed of migration.

### The Notch ligands Delta and Jagged are upregulated in winner GSCs

We have carried out both bulk mRNA sequencing and single cell quantitative RT-PCR on ALDH+ cells [10]. Although not a highly enriched population, differential gene expression analyses revealed a significant increase in components of the Notch pathway, including Delta and Jagged. Next, we directly assessed Delta, Jagged and Notch expression by qPCR from FACS isolated GSCs isolated from both winner and loser genotypes. We found that, on average, a Jagged 2 (JAG2) homolog was upregulated ca. 8X, and Delta 6X in winners vs loser GSCs (Figure 1C). In contrast, expression of Notch was equivalent in between both genotypes (Figure 1C).

The parallel increase in JAG2 expression, cluster size and migration kinetics in winner GSCs was intriguing, as a similar mechanism have been described in metastasis. In mammals, Jagged expression positively correlates with cancer invasiveness [22,23], as well as playing a role in collective migration [24]. In addition, multiple studies suggest that aggressiveness and clustering may be linked [25]. We next took a candidate approach, focusing on manipulating GSC behavior.

### HGF stimulation results in increased JAG2 expression, and a concurrent increase in migratory ability and cluster size in loser GSCs

It has been previously shown that several growth factors effect both stem cell migration as well as expression of proteins in the Notch pathway, and one interesting candidate was Hepatocyte Growth Factor (HGF). In several cancer models, HGF has been shown to increase both expression of jagged and tumor invasiveness [26–29], including human germ cell tumors [30]. In addition, HGF has been shown to play a role in collective migration of epithelial cells [31]. Finally, Botryllus GSCs express a HGF receptor (c-met) homolog. Thus, we next tested HGF stimulation on migration. Following dose dependency and viability assays (see methods) isolated GSCs were incubated in 10ng HGF and then tested for both JAG2 expression levels and migration to S1P.

HGF treatment of GSCs resulted in increased expression of JAG2 in both cell populations as seen in qRT-PCR (Figure 2A). Loser GSCs had increased JAG2 expression levels (p<0.001) and were indistinguishable from untreated winner GSCs, (p<0.001) (Figure 2A). Interestingly HGF-treated winner GSCs only had a slight increase in JAG2 expression, suggesting that there is either an upper threshold of Jagged expression, or of HGF stimulation.

**Figure 2:**
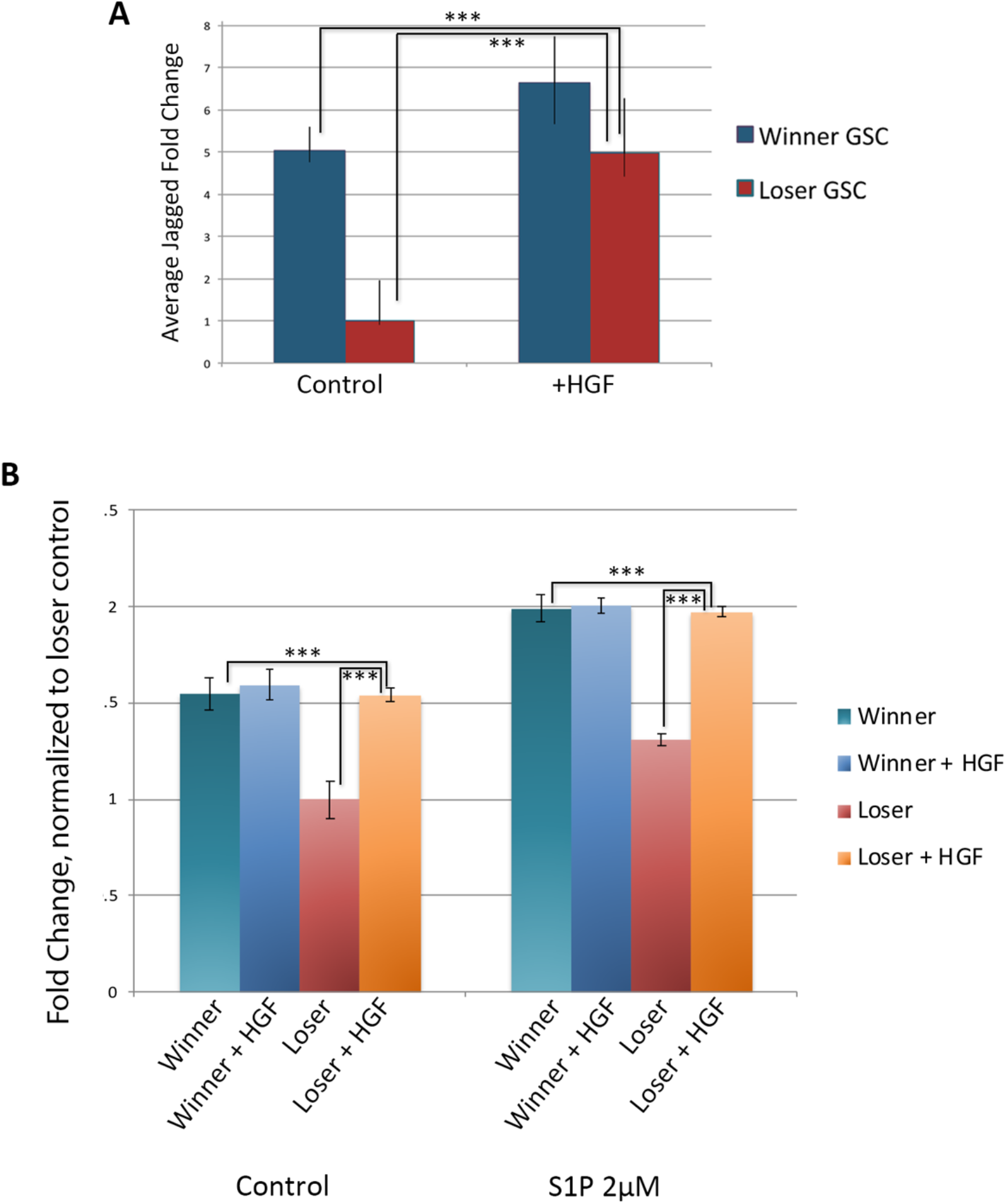
Treatment with Hepatocyte Growth Factor (HGF) increases jagged expression and migratory activity of loser GSCs. **A.** Quantification of jagged expression following treatment of winner and loser GSCs with HGF for 2 h. Data shown as relative fold change normalized to loser GSCs. ***p<0.001 (Students t-test). **B.** Results from transwell migration assays following HGF treatment on winner and loser GSCs. Show is the fold change of total cell count under different conditions, normalized to loser GSCs (n=6). ***p<0.001.

Transwell migration assays showed that loser GSCs had a 1.5X increase in migratory ability, both in control and in response S1P (p<0.001), essentially converting the loser GSC migration phenotype to that of the winner GSC phenotype. Interestingly, there was no statistical difference between winner GSC migration levels and HGF-treated loser GSCs (Figure 2B). Similar to the mild increase in expression of JAG2 following HGF treatment, winner GSCs treated with HGF had virtually no change in their migratory capabilities.

Analysis of the cell clustering revealed that HGF significantly increased clustering of the loser GSCs to a level equivalent to winner GSCs (Figure 5B). In contrast, there was no observable differences in cluster size of the winner GSCs between control vs. HGF (Figure 5). In summary, HGF treatment of loser GSCs caused them to take on a winner phenotype in three categories: JAG2 expression, migration kinetics and cluster size, but had little effect on winner genotypes.

### Perturbation of MAPK pathway results in changes of JAG2 expression, migratory ability and cluster size in winner GSCs

HGF is known to activate the MAPK and Akt pathways [26,28,29], both of which have been shown to be upstream of Jagged expression. In the next set of experiments, we used small molecule inhibitors to block both pathways. We utilized the MEK inhibitor U0126 and the AKT/P13 Kinase inhibitor Ly294002. Inhibiting the MAPK pathway using U0126 has been shown to result in decreased Jagged expression in several systems, for example squamous cell carcinoma cells, HUVEC, and breast cancer cell [32–34]. Ly294002 inhibits Akt through P13 Kinase [34] and has been shown to attenuate Jagged expression as well as mitigate activation of the Akt pathway following addition of HGF [35,36].

Following dose and viability assays (see methods), GSCs were incubated 10μM of U0126 for and assessed for Jagged expression. Jagged expression levels decreased up to three-fold in winner GSCs, with treated winner GSCs having similar expression levels to those of loser GSCs (Figure 3A). In contrast UO126 treatment only mildly decreased JAG2 expression. This suggests that, opposite to HGF stimulation, there may be a lower threshold of JAG2 expression, and/or jagged expression is also maintained by independent signals.

**Figure 3:**
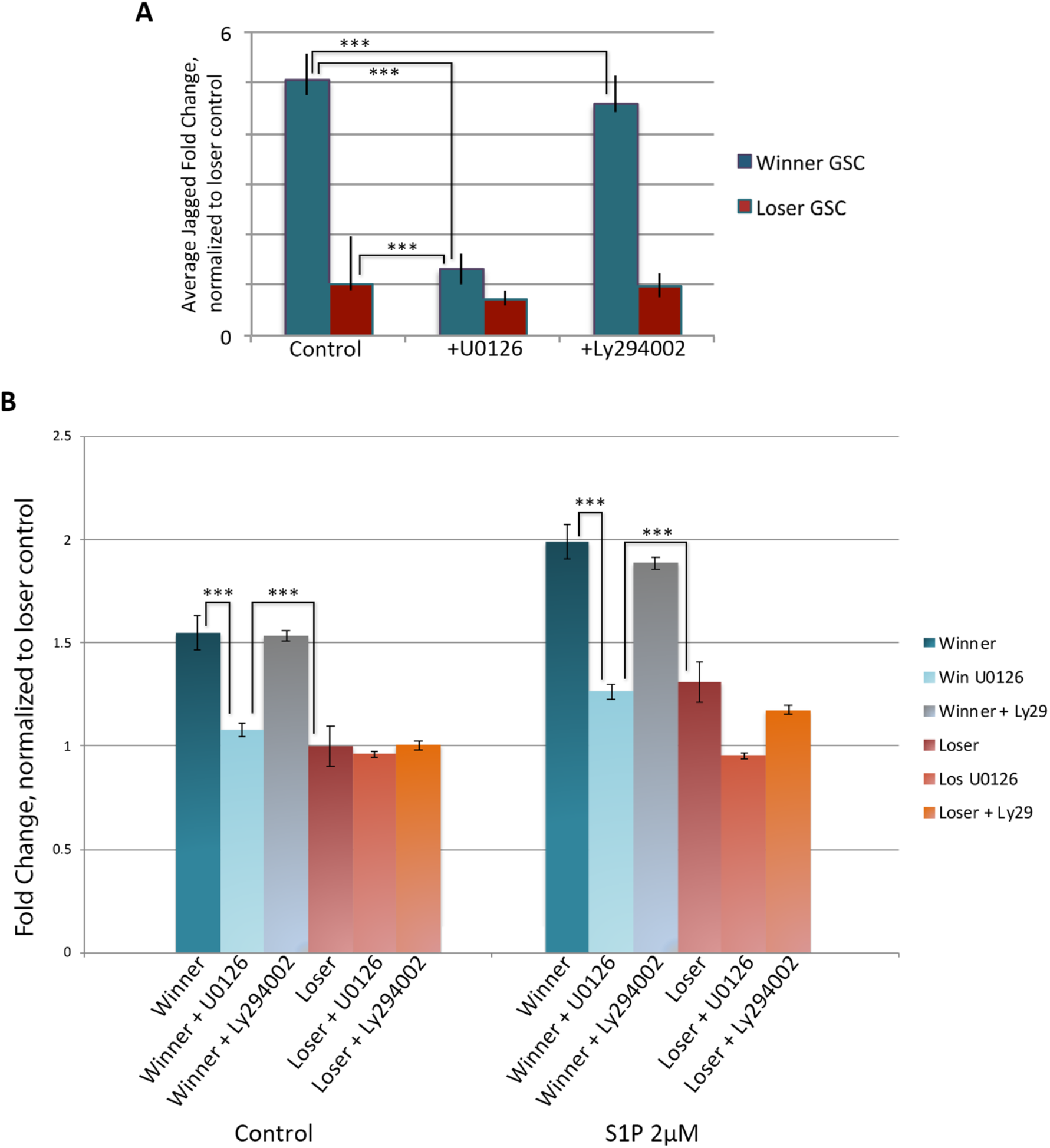
Blocking the MAPK pathway with UO126 decreases jagged expression and migratory ability. **A.** Treatment with UO126 results in decreased jagged expression in winner GSCs and has a small effect on loser GSCs as assessed by qPCR. Treatment with Ly294002 which blocks Akt activity had no effect. **B.** Transwell migration assays of GSCs following UO126 and Ly294002 treatment. Addition of UO126 to winner GSCs resulted in decreased migration to that nearly equivalent of loser GSCs under both control and S1P stimulated conditions. Loser GSCs showed a small effect from UO126 under S1P conditions. In contrast, Ly294002 had virtually no effect on winner or loser GSCs. Data expressed as averages of relative fold change, normalized to loser GSCs under control conditions. n=27. ***p<0.001.

Transwell migration assays revealed that MAPK inhibition with UO126 resulted in a decrease of the migration levels of the winner GSCs, decreasing them to the level of the loser GSCs (p<0.001; Figure 3B). In contrast, migration levels of the loser GSCs were not significantly lessened with MEK1/2 inhibition.

Analysis of the cluster size of the winner and loser GSCs following migration gave the exact opposite results of HGF (Figure 5 C, D). In this case, the winner clusters had decreased in size, with a distribution reminiscent of the untreated loser GSCs. In contrast, the loser GSCs were unaffected, again revealing a correlation between cluster size and migration kinetics.

We next inhibited Akt signaling using LY249002. Although Ly249002 has been used successfully in a number of ascidian species [37], it had no measurable effect on migration of either cell population, JAG2 expression levels, or cluster size, suggesting that the MAPK pathway and not the AKT pathway is involved in winner phenotype migration (Figure 3A, B).

To further validate the finding that the MAPK pathway specifically affects Jagged expression and influences migratory ability, we next used the selective ERK1/2 inhibitor FR180204 [38]. Inhibition with FR108204 resulted in a three-fold reduction in JAG2 expression, similarly to that of U0126 treatment as quantified with RT-PCR (Figure 4A; p<0.001). Equivalent to U0126 treatment, addition of FR108204 decreased migratory ability of winner GSCs, and winner GSC migration levels matched the level of untreated loser GSCs (Figure 4B; p< 0.001). Analysis of cluster size following migration showed equivalent results to UO126 treatment: winner clusters had decreased in size, while loser GSCs were unaffected by inhibition of MAPK signaling. Thus, both MAPK inhibitors showed equivalent results.

**Figure 4:**
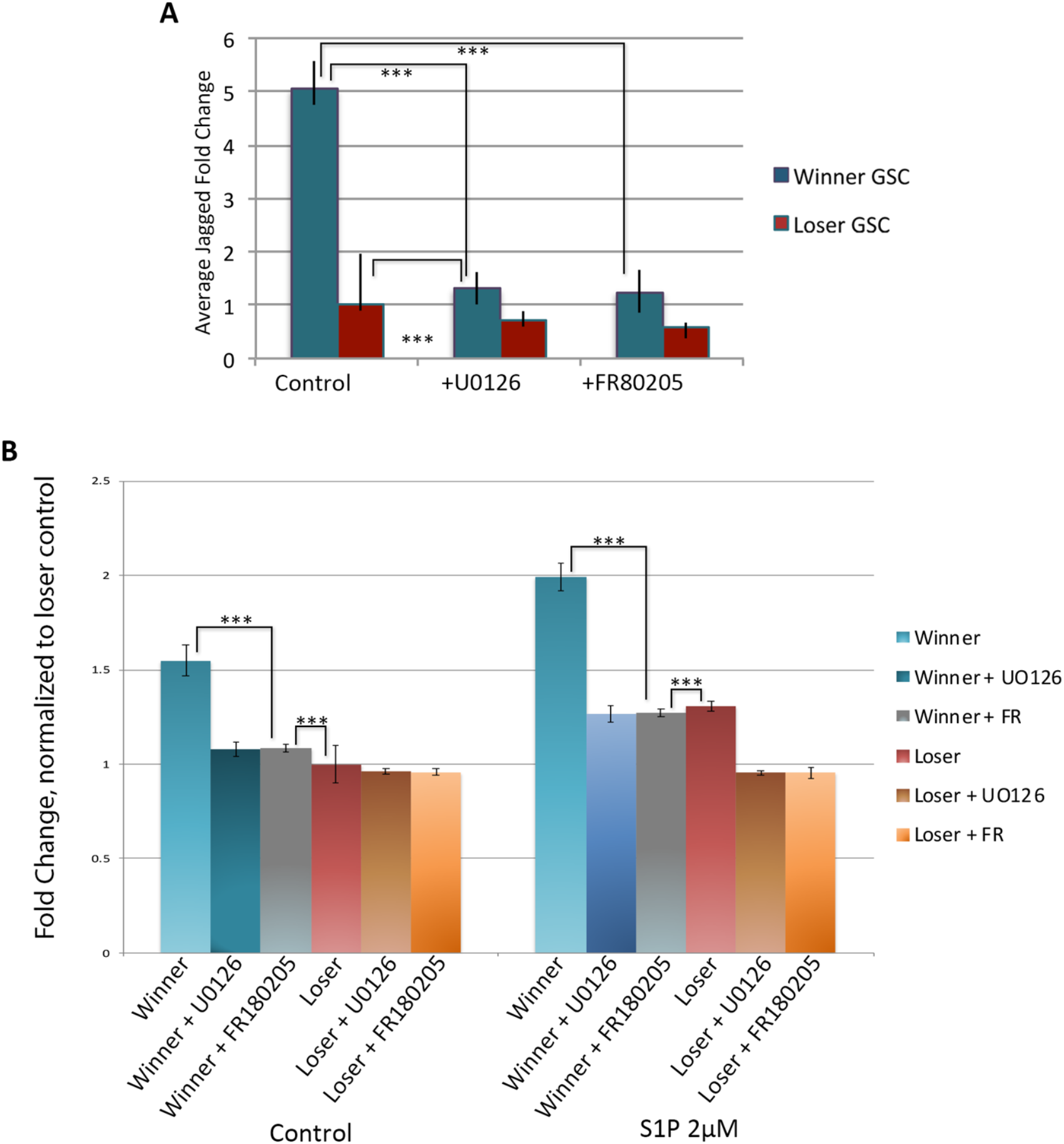
Blocking the MAPK pathway using FR180205 is equivalent to UO126 treatment. **A.** Treatment with both MAPK inhibitors had equivalent effects on jagged expression on winner, but not loser GSCs as quantified using qPCR. Data expressed as averages of relative fold change, normalized to loser GSCs. SDEVs were calculated for each average expression ratio (n=5) ***p<0.001 (Students t-test). **B.** Results from transwell migration assays following blocking MAPK activity using UO126 and FR180205 decreases migratory ability of winner GSCs but has a small effect on loser GSCs under both control and S1P stimulated conditions. Data expressed as average fold change in total cell count normalized to loser control. n=15. ***p<0.001.

**Figure 5:**
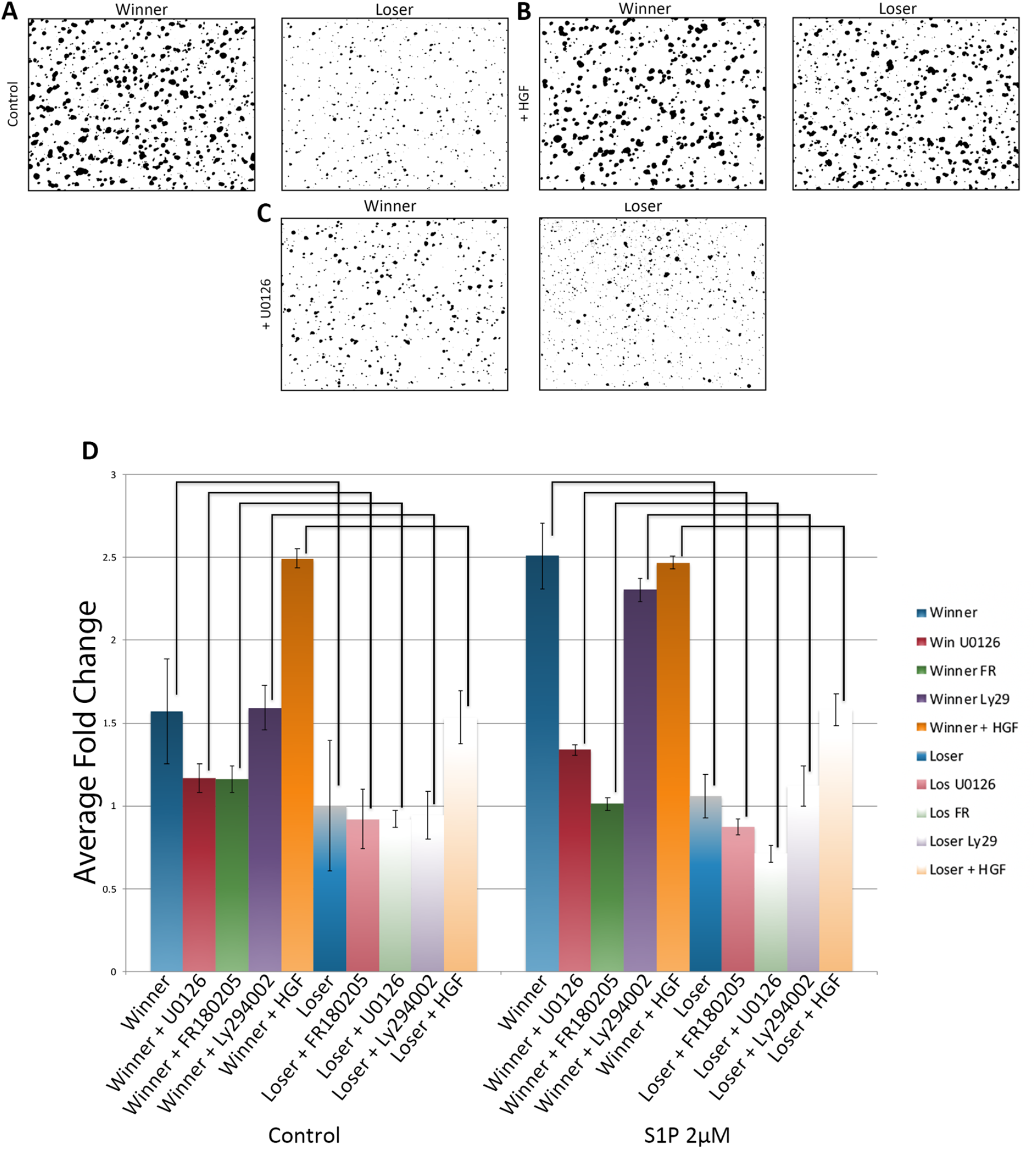
Genotype specific manipulation of cluster size. **A-C.** Representative images of cell clusters following transwell migration assays. **A.** under control conditions winner GSCs are found in larger clusters versus loser GSCs. **B.** Treatment with HGF causes clustering of loser GSCs but has no major effect on winner GSCs. **C.** Blocking MAPK signaling with UO126 has no effect on loser GSCs, but lowers cluster size in winner GSCs. **D.** Summary of average cell cluster size from all three winner loser pairs following HGF stimulation and MAPK inhibition. Data is normalized to the average cluster size of loser GSCs under control conditions. *p<0.05 **p<0.01 ***p<0.001. n=30 (control); n=27 (Ly294002 treatment); n=15 (UO126 and FR180205 treatments), n=9 (HGF treatment).

These data support a link between JAG2 expression via the MAPK pathway. When loser GSCs were stimulated with HGF their JAG2 expression levels matched those of untreated winner GSCs. Furthermore, following HGF treatment loser GSCs gained migratory function, to levels that phenocopied those of winner GSCs. On the flip side, inhibition of the MAPK pathway through either MEK1/2 (U0126) or ERK1/2 (FR180204) in winner GSCs resulted in not only a decrease of JAG2 expression, but also a decrease in migratory ability and cluster size. In all cases, Jagged expression, migration kinetics and cluster size responses to these treatments are linked.

These data also suggest that regulation of JAG2 expression is very specific, with a tightly controlled minimal and maximum level. Winner GSCs had very little response to HGF treatment, either in actual expression levels of JAG2 or in migratory abilities. Conversely, inhibition of the MAPK pathway through either inhibition of MEK1/2 or ERK1/2 resulted in a minimal decrease in either JAG2 expression or migratory levels of loser GSCs.

### Live imaging of the GSC niche following transplantation of winner and loser GSCs

Next, we assessed the behavior of FACS-isolated labeled winner and loser GSCs in vivo using live imaging. Isolated winner GSCs were labeled with CellTracker CM-DiI Dye and loser GSCs were labeled with Syto59. Labeled cells were then co-injected into the loser genotype at stage just prior to the normal GSC migration period between the old and new niche, and 24 h later, when GSC migration is maximal, we imaged the new germline niche of the recipient colony using spinning disc confocal microscopy (Figure 6A-I; the niche immediately inside the protruding region of the white dotted line [11]). In all three winner/loser pairs, we observed labeled cells in ca. 70% of the niches observed. In each positive niche, between 1-5 cells were found, with a slight increase in the number of winner GSCs versus losers (1.9X; p=0.018; Figure 6J). This was a minority of the cells that were injected, most of which remained in circulation and were moving too quickly to image. In addition, labeled cells that had migrated to the new niche remained stationary during the observation window.

**Figure 6:**
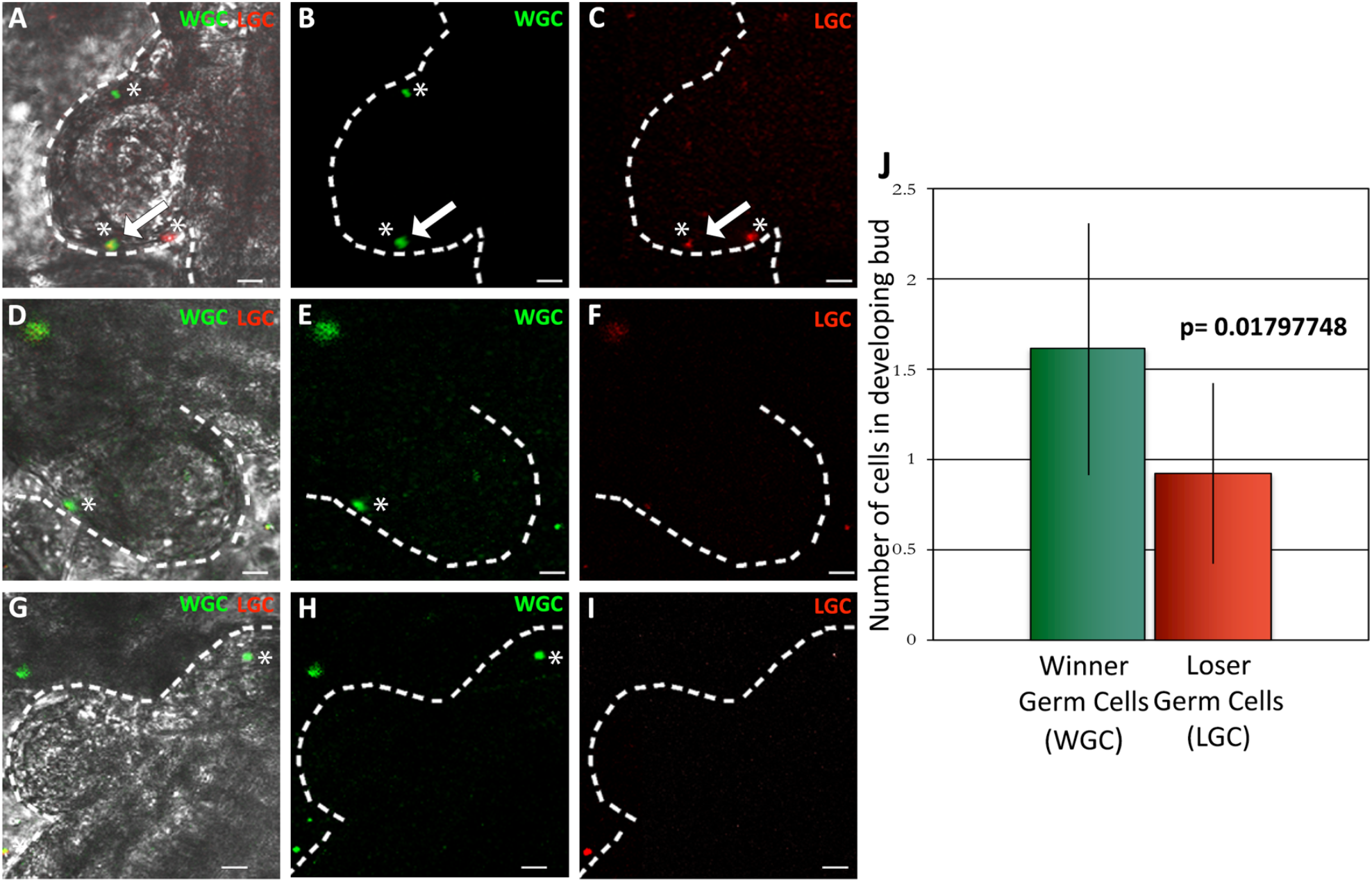
Live imaging of the GSC niche following transplantation of fluorescently labeled winner and loser GSCs. GSCs from each winner and loser pair were isolated by FACS and labeled with cell tracker dyes (winner GSCs (WG green); loser GSCs (LG red)) and co-injected into a subclone of the loser genotype. 24 h later the newly developing GSC niche was imaged (**A-I)**. The dotted white line is superimposed on the epidermis, the evagination is the newly developing zooid, and the niche is between the white line and the developing zooid viscera (see Supplemental Figure 1). Representative images of the three (A-C; D-F; G-I) types of phenotypes observed in this experiment, described in the text. Superimposed images of light and both fluorescent channels are shown in panels A, D, and G. **J.** Labeled cells were observed in ca. 70% of the GSC niches visualized, shown is the quantification of the number of winner vs loser GSCs observed (n=9). Winner GSCs had a slight advantage versus loser GSCs (1.9X; p<0.02).

Representative images of the range of distribution of the labeled cells we observed are shown in Figure 6. On average, about half the labeled cells observed in these experiments were found in pairs (Figure 6A, D), and there were no noticeable differences in these results between any of the winner/loser genotypes. We observed pairs of winners (top asterisk, Fig 6 A-E) and also pairs consisting of winners and losers (bottom asterisk, Fig 6 A-C). Single cells were also observed (Fig 6C far right; 6H). Unfortunately, we did not visualize a migration event in any experiment, but only observed cells once they had settled in the region being observed. In summary, in the time span between FACS isolation of single cells and imaging of the niche we do observe pairing of about half the labeled cells, but do not know if cells migrated as pairs, or interacted following arrival at the niche.

## Discussion

Here we utilized the natural phenotypic variation in germ cell parasitism to test the hypothesis that differences in S1P-induced migration contribute to the competitive abilities of GSCs. We found GSCs retained their competitive phenotype, initially characterized following in vivo interactions between different genotypes, when assessed ex vivo. Specifically, GSCs isolated from winner colonies migrated along a gradient of S1P faster than those from losers. We also found that this increase in migratory ability correlated to an increase in both the expression of the notch ligand, JAG2, as well as size of the cell clusters. JAG2 expression, clustering and migration speed could be increased by stimulation with the growth factor HGF, and decreased via inhibition of the MAPK pathway. But unexpectedly, these affects were genotype specific: HGF treatment converted GSC losers to a winner phenotype, while MAPK pathway inhibition converted winner GSCs to a loser phenotype. This suggests that collective migration may be part of germline stem cell competition, and that there may be minimum and maximum thresholds for all three of these phenotypes. Finally, live imaging of germline niches following transplantation of winner and loser cells revealed a slight, but significant increase winner cell occupancy, as well as some small clusters of cells. However, during these experiments we did not capture images of cells arriving in the niche, only after they had settled, thus do not know if these interactions occurred prior to or during migration, or occurred on the niche.

Increased collective migration by winner GSCs may allow more efficient and faster migration to the newly developing niche. This is likely due to specialization within the cluster, leading to formation of leader and follower cells (reviewed in [39,40]). Leader cells extend filopodia and lamellipodia, remodel the extracellular matrix and sense the microenvironment, providing direction for the follower cells [39], while follower cells may generate the dominant traction forces for migration [41]. Interestingly, in a study on human lung cancer metastatic lesions, leader and follower cells were separated and each population characterized. Leader cells showed increased expression of Jagged-1, Myosin-X, and fibronectin, which together affect the geometry of the leader cell filopodia and 3D invasion potential of the metastatic clusters [23]. Interestingly, the top 5 genes found to be upregulated in leader cells in that study are also upregulated in winner GSCs from Botryllus, including: Jagged, Myosin-X, fibronectin, RIP4K and ALR4C [23]. In addition, a recent study in collective migration of MDCK cells has shown that specification of leaders includes a positive feedback loop between HGF and ERK signaling [31]. Together, this suggests that Botryllus uses conserved mechanisms to increase collective migration in winner GSCs, perhaps by priming winner GSCs for leader cell fate, and further that these can be manipulated in vitro by stimulating or inhibiting the MAPK pathway. Besides being faster at migration, larger clusters may also be able to overpower smaller clusters, and push them off the niche.

Jagged expression has also been shown to be involved in a metastatic process which results in the formation of a hybrid epithelial/mesenchymal phenotype that correlates with collective migration [24]. Unfortunately, one issue we have not resolved yet is if there is if JAG2 expression is correlative with, or causative of changes cluster size, as experiments blocking notch signaling using DAPT both in vitro and in vivo were not consistent, despite the fact that we have used these inhibitors successfully before [42]. Dissecting a causal link between Notch signaling and cluster size is currently an active area of research in the lab. Similarly, while the Botryllus GSCs express a conserved HGF receptor (c-met) protein, there is not a clear homolog of HGF. In summary, while the ability to simultaneously manipulate JAG2 expression, cluster size and migration speed are robust, at this point the underlying molecular mechanisms are unknown.

The potential for clustering during GSC migration and its role in competition may be due to another aspect of chemotaxis in Botryllus, the generation of autonomous gradients, a phenomenon known as relay signaling [43,44]. We recently found that Botryllus GSCs may require an autonomously generated chemotactic gradient of the bioactive lipid 12-S-HETE for migration along an S1P gradient [21]. It is becoming appreciated that chemotaxis may require the migrating cell to contribute spatial information via creating autonomous gradients [45–47]. In turn, these autonomously-generated gradients cause self-organization of the cells, such that they migrate as a group [46,47]. Although we have not detected differences in mRNA expression of the 12-S-HETE biosynthetic proteins, or the 12-S-HETE receptor (GPR31) between winner and loser GSCs, in human prostate cancer expression levels of GPR31 and 12-S-HETE both positively correlate with metastasis [48,49]. The link between relay signaling, clustering and competition is the subject of current studies.

Differences in GSC cluster sizes observed ex vivo in this study are consistent with previous studies from our lab done on germline migration in vivo-with one major difference [11]. Using whole mount in situ hybridization, we had previously found that GSCs, defined by the expression of germline markers (e.g., vasa/piwi), always migrate in clusters between niches. Moreover, a large range of cluster sizes was observed in this study-while over half were made up of 10 cells or less, many larger groups were observed, including some with more than 100 cells. This is exciting, as these results were from observations of 241 GSC clusters in 21 animals, but was done without regard to gcp phenotype of the individuals tested [11]. However, while these results clearly showed that GSCs migrate in clusters, and that the sizes can be widely variable, they also revealed that the GSC clusters always included another cell type. This second cell does not express vasa or piwi, but does express a TGFβ family member (TGFβ-f; [11,20]). In addition, the two cell types were associated with each other over 99% of the time. Moreover, these vasa+/ TGFβ-f+ clusters are found in every stage of adulthood, from the oozooid, to the fertile adult [11].

These results bring up an issue we do not yet understand-the lineage relationship between the FACS isolated, single IA6+ cells studied here and previously [20,21], and the clustered vasa+ cells detected by in situ hybridization [11]. First, all of our functional studies have been based on enrichment of adult progenitor cells by FACS using various methods, including levels of ALDH enzymatic activity, cell surface lectins, and an antibody to IA6 [10,20,21]. These are single cells that are highly enriched in expression of germline markers, including vasa and piwi, can be transplanted and will give rise to mature gametes [20]. However, in situ hybridization of both vasa and IA6 identifies clusters of cells in the germline niche in fertile adults [11,20]. In addition, there is a size difference in the FACS isolated IA6+ cells, which are in the 6-10 um range [10,20] - and the GSCs identified by in situ hybridization using vasa or IA6, which are larger, in the 8-12 um range [11,20]. In summary, at this point we do not know the relationship between the single IA6+ cells isolated by FACS, and the vasa+ cells found associated with TGFβ-f+ cells in clusters observed in vivo. However, it is clear that isolated IA6+ cells can cluster within a few hours of dissociation both in vitro and in vivo, and also migrate to the niche following transplantation (Figure 6).

One observation suggests that IA6+ cells may be a precursor to both the germline progenitor and TGFβ-f+ cell. When GSCs are transplanted between genetically distinguishable individuals, isolated testis and oocytes are clonal, and either derived from the donor or the recipient genotype [9,10]. TGFβ-f+ cells remain associated with mature gametes, TGFβ-f is highly expressed in follicle cells that encompass both mature gametes [11], and these cells are not usually removed during isolation of mature gametes [10,15,17]. This suggests that IA6+ cells isolated by FACS are precursors to both the GSCs-defined by expression of germline markers; and associated TGFβ-f+ expressing somatic niche cells-also defined by the absence of expression of germline markers. In summary, while we clearly can see migration of labeled IA6+ cells to the niche in vivo (Figure 6 and [20]), we do not yet know if the cells isolated by FACS and tested here are from dissociated clusters, or another population not identified by in situ hybridization. We hypothesize it is the former, as clusters of vasa+/ TGFβ-f+ cells are observed in the oozooid, immediately following metamorphosis, and are found in the colony in every stage of development [20]. And while single cells expressing vasa, IA6 or TGFβ-f have been observed, they are at very low frequency, < 1% of the cells [11,20]. Together, this suggests that single IA6+ progenitors are from clusters dissociated during isolation and have the ability to proliferate and differentiate to reform vasa+/ TGFβ-f+ clusters. This is consistent with previous limit dilution studies [20], in which a small number of transplanted cells (n=5) gave rise to multiple testis in the recipient, and this chimerism was maintained for months.

Live imaging experiments revealed both a slight increase in winner GSCs vs. losers in individual niches, as well as interactions between winner and loser cells. However, as we did not observe a migration event, we could not discriminate between collective migration versus interactions occurring after arrival of cells to the niche. As the process takes multiple days, further imaging between 12-72 hours post injection could identify key timeframes of niche entry, as well as any potential interactions that occur on the niche.

Previous studies using situ hybridization found large clusters of GSCs in the new niche at the exact same timepoint which was being imaged here [11], so we were initially disappointed at the low number of cells observed in vivo. However, an important potential confounding factor to these results is that the niche itself may be limiting. While the location of the new niche is known [11], its structure is not clear. EM studies reveal it is a transparent region between the bud epithelium and the epidermis (immediately inside the white dotted line in Figure 6) that seems to be made of extracellular matrix, and GSCs do not seem to be associated with either cell layer. (Di Maio, unpublished). Thus, it is important to note that in these transplantation experiments the recipient is unmanipulated, and has a normal level of unlabeled GSCs that are also present and migrating. This is unlike other experimental systems, for example, those on mammalian hematopoietic stem cell function, where the recipient stem cell niches are cleared via radiation or chemical treatments prior to transplantation. This is interesting as we found that the labeled IA6+ cells which did not migrate and settle in the new niche remained in circulation. Cells also remained in circulation when sphingosine-1-phosphate receptor (S1PR1) signaling was blocked in a previous study, which prevented GSCs from migrating to the niche [20]. In summary, the presence of unlabeled donor cells in the niche may explain our observations. Importantly, donor derived gametes are not usually assessed until weeks following transplantation, regardless if this occurred naturally via vascular fusion, or microinjection of FACS isolated progenitors [9,10,13–17]. Thus, the 1.9X increase in migration by winner genotypes, as well as identification of small clusters of labeled cells appearing in the niche of an unmanipulated recipient 24 h following transplantation, may be more significant than they appear.

Another possible explanation is based on the fact that GSCs migrate from the old niche to the new niche in a synchronized fashion that is linked to somatic development, and is controlled in part by secretion of S1P from the new GSC niche [11,20]. In these experiments, GSCs were not isolated with any regard to the stage of asexual development and it would be no surprise if stage specific gene expression patterns controlled clustering and migratory ability. Conversely, we do not see major changes in jagged or chemotactic genes (e.g., S1PR1) in stage specific transcriptome data [50], nor was there any effect of stage on any of the in vitro assays here or in previous studies [20,21]. Finally, cells are being dissociated, treated with antibodies, sorted by FACS, and microinjected, and this introduces multiple potential artifacts-although this would not be unique to Botryllus or this study.

In summary, here we have found that winner GSCs show enhanced migratory ability to chemotactic cues ex vivo, and that enhanced migration correlates with both expression of the notch ligand, jagged, as well as cluster size. Collective migration and a plasticity to cluster size is completely consistent with previous studies done in vivo, which found over a 10X range of cluster sizes among different genotypes [20]. While we were unable to show that cluster size was due to notch signaling, the Notch pathway has been implicated in clustering and leader/follower differentiation in a number of cell types, during both normal and metastatic collective migration [22,24,51–53]. In addition, the role of HGF in specification of leader cells has also been demonstrated [31], and the original description of HGF was based on an induced ‘scatter’ phenotype in MDCK cells that is remarkably similar to results here [54]. This suggests that Botryllus GSCs use conserved mechanisms during cell migration and that these are likely part of the cellular and molecular underpinnings of germ cell competition. The ability to study conserved aspects of cell migration both in vivo and in vitro, coupled to the genotypic variation provided by the competitive properties of GSCs, make Botryllus an excellent model for future studies on competition, chemotaxis and collective cell migration.

## Acknowledgements

We would like to thank Greg Stoney for his outstanding care of the mariculture facility, and Susannah Kassmer, Delany Rodriguez and Shane Nourizadeh for all their guidance and help.

## Author Contributions

MF and AD conceived and designed the experiments, MF conducted the experiments, and MF and AD wrote the manuscript.

---

**Supplemental Fig 1.**
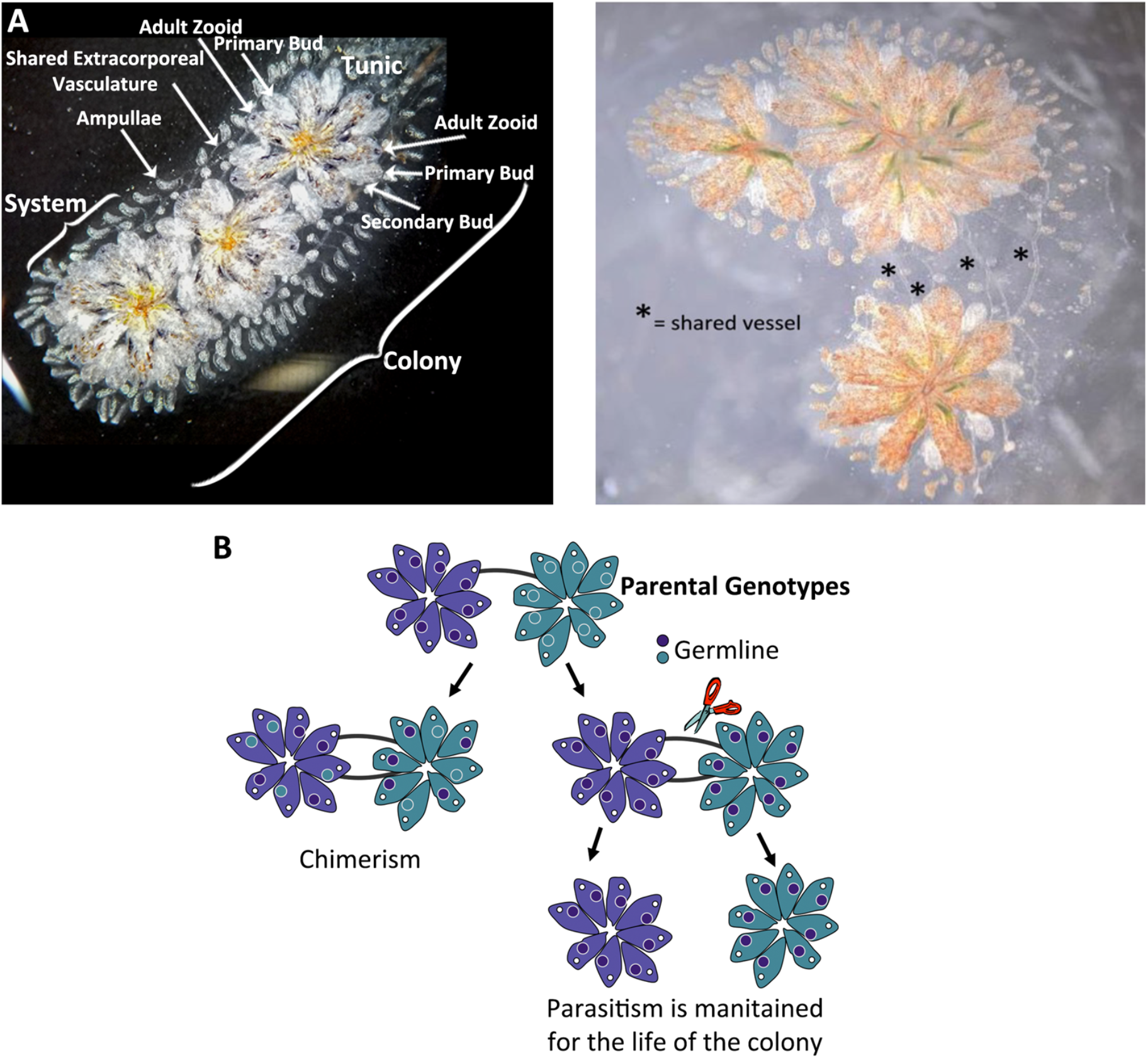
*Botryllus schlosseri* morphology and Germline Stem Cell Parasitism. **(A)** Ventral view of a *Botryllus schlosseri* colony. The colony is the combination of several rosette shaped systems composed of adult animals (zooids). Individual animals are connected by a shared each of extracorporeal vasculature, which ends in bulb like extensions called ampullae. Each zooid has asexually developing primary buds and secondary buds. During the asexual budding process. As new buds form during a weekly regeneration process germ cells (germline stem cells (GSP) migrate into the developing buds via shared vasculature (***) to the GSP niche. **(B)** Germ cells destined for the developing secondary bud germline niche migrate through the extracorporeal vasculature and contribute to the developing bud. Fusion of related individuals results in each genotypes mobile progenitors moving between animals. GSCs of two individuals are mixed, one genotype will outcompete the other, solely contributing to the germline of subsequent generattons. Termed stem cell parasitism (SCP) this clonal dominance is heritable, stable and repeatable.

**Supplementary Fig 2.**
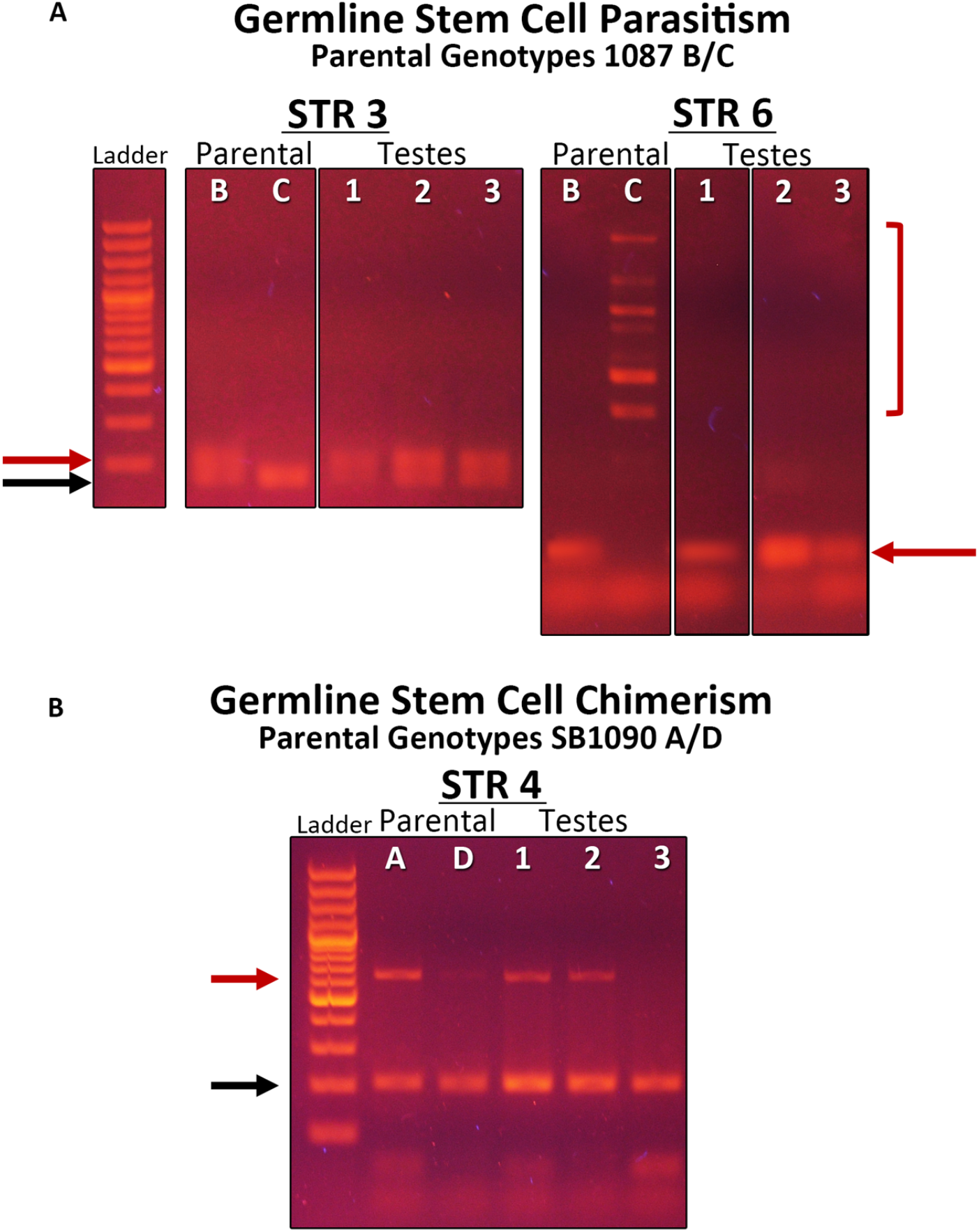
Examples of parasitism and chimerism detected using microsatellite (STR) PCR. (A) Example of germline stem cell parasitism following fusion of genotypes 1087B and 1087G detected using two loci, STR3 (codominant marker) and STR6 (dominant marker) amplified from DNA of individual testis isolated 4 weeks following vascular fusion. Parental DNA samples are from somatic tissue isolated from naive subclones of the same genotype. In this case, both loci show that all testis in the chimera are derived from genotype 1087B. 1087B and 1087G are a winner/loser pair (B) Example of germline chimerism following fusion of genotypes 1090A and 1090D using STR4 (codominant marker). Individual testis in the chimera are derived from one genotype or the other. In this case testis from both genotypes are found throughout the chimera. Note that we have previously reported that testis are clonally derived [8,9], Black arrows are shared alleles at each STR loci, while red arrows and outline are polymorphic alleles

## References

1. Morata G, Ripoll P. Minutes: Mutants of Drosophila autonomously affecting cell division rate. Dev Biol. 1975;42: 211–221. doi:10.1016/0012-1606(75)90330-9

2. Amoyel M, Bach EA. Cell competition: how to eliminate your neighbours. Development. 2014;141: 988–1000. doi:10.1242/dev.079129

3. Slaidina M, Lehmann R. Quantitative Differences in a Single Maternal Factor Determine Survival Probabilities among Drosophila Germ Cells. Curr Biol. 2017;27: 291–297. doi:10.1016/j.cub.2016.11.048

4. Jin Z, Kirilly D, Weng C, Kawase E, Song X, Smith S, et al. Differentiation-Defective Stem Cells Outcompete Normal Stem Cells for Niche Occupancy in the Drosophila Ovary. Cell Stem Cell. 2008;2: 39–49. doi:10.1016/j.stem.2007.10.021

5. Bondar T, Medzhitov R. p53-Mediated Hematopoietic Stem and Progenitor Cell Competition. Cell Stem Cell. 2010;6: 309–322. doi:10.1016/j.stem.2010.03.002

6. Baker NE. Emerging mechanisms of cell competition. Nat Rev Genet. 2020;21: 683–697. doi:10.1038/s41576-020-0262-8

7. Herrera SC, Bach EA. Super-Competitors Game the Fitness Sensing System. Dev Cell. 2018;46: 672–674. doi:10.1016/j.devcel.2018.09.006

8. Delsuc F, Philippe H, Tsagkogeorga G, Simion P, Tilak M-K, Turon X, et al. A phylogenomic framework and timescale for comparative studies of tunicates. Bmc Biol. 2018;16: 39. doi:10.1186/s12915-018-0499-2

9. Brown FD, Tiozzo S, Roux MM, Ishizuka K, Swalla BJ, De Tomaso AW. Early lineage specification of long-lived germline precursors in the colonial ascidian Botryllus schlosseri. Development. 2009;136: 3485–3494. doi:10.1242/dev.037754

10. Laird DJ, De Tomaso AW, Weissman IL. Stem Cells Are Units of Natural Selection in a Colonial Ascidian. Cell. 2005;123: 1351–1360. doi:10.1016/j.cell.2005.10.026

11. Langenbacher AD, De Tomaso AW. Temporally and spatially dynamic germ cell niches in Botryllus schlosseri revealed by expression of a TGF-beta family ligand and vasa. Evodevo. 2016;7: 9. doi:10.1186/s13227-016-0047-5

12. Sabbadin A. Le basi genetiche della capacita di fusione fra colonies in Botryllus schlosseri (Asidiacea). Rend Accad Naz Lincie, Series 8. 1962;32: 1031–1035.

13. Sabbadin A, Zaniolo G. Sexual differentiation and germ cell transfer in the colonial ascidian Botryllus schlosseri. J Exp Zool. 1979;207: 289–304. doi:10.1002/jez.1402070212

14. Stoner DS, Rinkevich B, Weissman IL. Heritable germ and somatic cell lineage competitions in chimeric colonial protochordates. Proc National Acad Sci. 1999;96: 9148–9153. doi:10.1073/pnas.96.16.9148

15. Stoner DS, Weissman IL. Somatic and germ cell parasitism in a colonial ascidian: Possible role for a highly polymorphic allorecognition system. Proc National Acad Sci. 1996;93: 15254– 15259. doi:10.1073/pnas.93.26.15254

16. Pancer Z, Gershon H, Rinkevich B. Coexistence and Possible Parasitism of Somatic and Germ Cell Lines in Chimeras of the Colonial Urochordate Botryllus schlosseri. Biol Bull. 1995;189: 106–112. doi:10.2307/1542460

17. Carpenter MA, Powell JH, Ishizuka KJ, Palmeri KJ, Rendulic S, De Tomaso AW. Growth and Long-Term Somatic and Germline Chimerism Following Fusion of Juvenile Botryllus schlosseri. Biological Bulletin. 2011;220: 57–70. doi:10.1086/bblv220n1p57

18. Rosental B, Kowarsky M, Seita J, Corey DM, Ishizuka KJ, Palmeri KJ, et al. Complex mammalian-like haematopoietic system found in a colonial chordate. Nature. 2018;564: 425–429. doi:10.1038/s41586-018-0783-x

19. Braden BP, Taketa DA, Pierce JD, Kassmer S, Lewis DD, De Tomaso AW. Vascular Regeneration in a Basal Chordate Is Due to the Presence of Immobile, Bi-Functional Cells. Plos One. 2014;9: e95460. doi:10.1371/journal.pone.0095460

20. Kassmer SH, Rodriguez D, Langenbacher AD, Bui C, De Tomaso AW. Migration of germline progenitor cells is directed by sphingosine-1-phosphate signalling in a basal chordate. Nat Commun. 2015;6: 8565. doi:10.1038/ncomms9565

21. Kassmer SH, Rodriguez D, De Tomaso AW. Evidence that ABC transporter-mediated autocrine export of an eicosanoid signaling molecule enhances germ cell chemotaxis in the colonial tunicate Botryllus schlosseri. Development. 2020;147: dev184663. doi:10.1242/dev.184663

22. He W, Chan CML, Wong SCC, Au TCC, Ho WS, Chan AKC, et al. Jagged 2 silencing inhibits motility and invasiveness of colorectal cancer cell lines. Oncol Lett. 2016;12: 5193– 5198. doi:10.3892/ol.2016.5321

23. Summerbell ER, Mouw JK, Bell JSK, Knippler CM, Pedro B, Arnst JL, et al. Epigenetically heterogeneous tumor cells direct collective invasion through filopodia-driven fibronectin micropatterning. Sci Adv. 2020;6: eaaz6197. doi:10.1126/sciadv.aaz6197

24. Boareto M, Jolly MK, Goldman A, Pietilä M, Mani SA, Sengupta S, et al. Notch-Jagged signalling can give rise to clusters of cells exhibiting a hybrid epithelial/mesenchymal phenotype. J Roy Soc Interface. 2016;13: 20151106. doi:10.1098/rsif.2015.1106

25. Friedl P, Gilmour D. Collective cell migration in morphogenesis, regeneration and cancer. Nat Rev Mol Cell Bio. 2009;10: 445–457. doi:10.1038/nrm2720

26. Huang K-H, Sung I-C, Fang W-L, Chi C-W, Yeh T-S, Lee H-C, et al. Correlation between HGF/c-Met and Notch1 signaling pathways in human gastric cancer cells. Oncol Rep. 2018;40: 294–302. doi:10.3892/or.2018.6447

27. Tsou H-K, Chen H-T, Hung Y-H, Chang C-H, Li T-M, Fong Y-C, et al. HGF and c-Met interaction promotes migration in human chondrosarcoma cells. PLoS One. 2013;8: e53974. doi:10.1371/journal.pone.0053974

28. Zeng Q, Li S, Chepeha DB, Giordano TJ, Li J, Zhang H, et al. Crosstalk between tumor and endothelial cells promotes tumor angiogenesis by MAPK activation of Notch signaling. Cancer Cell. 2005;8: 13–23. doi:10.1016/j.ccr.2005.06.004

29. Viticchiè G, Muller PAJ. c-Met and Other Cell Surface Molecules: Interaction, Activation and Functional Consequences. Biomedicines. 2015;3: 46–70. doi:10.3390/biomedicines3010046

30. Scheri KC, Leonetti E, Laino L, Gigantino V, Gesualdi L, Grammatico P, et al. c-MET receptor as potential biomarker and target molecule for malignant testicular germ cell tumors. Oncotarget. 2018;9: 31842–31860. doi:10.18632/oncotarget.25867

31. Hino N, Matsuda K, Jikko Y, Maryu G, Sakai K, Imamura R, et al. A feedback loop between lamellipodial extension and HGF-ERK signaling specifies leader cells during collective cell migration. Dev Cell. 2022;57: 2290–2304.e7. doi:10.1016/j.devcel.2022.09.003

32. Sierra RA, Trillo-Tinoco J, Mohamed E, Yu L, Achyut BR, Arbab A, et al. Anti-Jagged Immunotherapy Inhibits MDSCs and Overcomes Tumor-Induced Tolerance. Cancer Res. 2017;77: 5628–5638. doi:10.1158/0008-5472.can-17-0357

33. Kiec-Wilk B, Grzybowska-Galuszka J, Polus A, Pryjma J, Knapp A, Kristiansen K. The MAPK-dependent regulation of the Jagged/Notch gene expression by VEGF, bFGF or PPAR gamma mediated angiogenesis in HUVEC. J Physiol Pharmacol. 2010;61: 217–25.

34. Hu L, Zaloudek C, Mills GB, Gray J, Jaffe RB. In vivo and in vitro ovarian carcinoma growth inhibition by a phosphatidylinositol 3-kinase inhibitor (LY294002). Clin Cancer Res. 2000;6: 880–6.

35. Roux PP, Blenis J. ERK and p38 MAPK-activated protein kinases: a family of protein kinases with diverse biological functions. Microbiol Mol Biol Rev. 2004;68: 320–44. doi:10.1128/mmbr.68.2.320-344.2004

36. You B, Yang Y-L, Xu Z, Dai Y, Liu S, Mao J-H, et al. Inhibition of ERK1/2 down-regulates the Hippo/YAP signaling pathway in human NSCLC cells. Oncotarget. 2015;6: 4357–68. doi:10.18632/oncotarget.2974

37. Lambert CC. Signaling pathways in ascidian oocyte maturation: effects of various inhibitors and activators on germinal vesicle breakdown. Dev Growth Differ. 2005;47: 265–72. doi:10.1111/j.1440-169x.2005.00796.x

38. Ohori M, Takeuchi M, Maruki R, Nakajima H, Miyake H. FR180204, a novel and selective inhibitor of extracellular signal-regulated kinase, ameliorates collagen-induced arthritis in mice. Naunyn Schmiedebergs Arch Pharmacol. 2007;374: 311–6. doi:10.1007/s00210-006-0117-7

39. Mayor R, Etienne-Manneville S. The front and rear of collective cell migration. Nat Rev Mol Cell Bio. 2016;17: 97–109. doi:10.1038/nrm.2015.14

40. Mercedes SAV, Bocci F, Levine H, Onuchic JN, Jolly MK, Wong PK. Decoding leader cells in collective cancer invasion. Nat Rev Cancer. 2021;21: 592–604. doi:10.1038/s41568-021-00376-8

41. Trepat X, Wasserman MR, Angelini TE, Millet E, Weitz DA, Butler JP, et al. Physical forces during collective cell migration. Nat Phys. 2009;5: 426–430. doi:10.1038/nphys1269

42. Kassmer SH, Langenbacher AD, De Tomaso AW. Integrin-alpha-6+ Candidate stem cells are responsible for whole body regeneration in the invertebrate chordate Botrylloides diegensis. Nat Commun. 2020;11: 4435. doi:10.1038/s41467-020-18288-w

43. Garcia GL, Parent CA. Signal relay during chemotaxis. J Microsc. 2008;231: 529–34. doi:10.1111/j.1365-2818.2008.02066.x

44. SenGupta S, Parent CA, Bear JE. The principles of directed cell migration. Nat Rev Mol Cell Bio. 2021;22: 529–547. doi:10.1038/s41580-021-00366-6

45. Insall RH. Receptors, enzymes and self-attraction as autocrine generators and amplifiers of chemotaxis and cell steering. Curr Opin Cell Biol. 2023;81: 102169. doi:10.1016/j.ceb.2023.102169

46. Insall RH, Paschke P, Tweedy L. Steering yourself by the bootstraps: how cells create their own gradients for chemotaxis. Trends Cell Biol. 2022;32: 585–596. doi:10.1016/j.tcb.2022.02.007

47. Stock J, Pauli A. Self-organized cell migration across scales – from single cell movement to tissue formation. Development. 2021;148: dev191767. doi:10.1242/dev.191767

48. Honn KV, Guo Y, Cai Y, Lee M-J, Dyson G, Zhang W, et al. 12-HETER1/GPR31, a high-affinity 12(S)-hydroxyeicosatetraenoic acid receptor, is significantly up-regulated in prostate cancer and plays a critical role in prostate cancer progression. FASEB J. 2016;30: 2360–9. doi:10.1096/fj.201500076

49. Dilly A, Tang K, Guo Y, Joshi S, Ekambaram P, Maddipati KR, et al. Convergence of eicosanoid and integrin biology: Role of Src in 12-LOX activation. Exp Cell Res. 2017;351: 1–10. doi:10.1016/j.yexcr.2016.12.011

50. Rodriguez D, Sanders EN, Farell K, Langenbacher AD, Taketa DA, Hopper M, et al. Analysis of the basal chordate Botryllus schlosseri reveals a set of genes associated with fertility. Bmc Genomics. 2014;15: 1183. doi:10.1186/1471-2164-15-1183

51. Boareto M, Jolly MK, Lu M, Onuchic JN, Clementi C, Ben-Jacob E. Jagged–Delta asymmetry in Notch signaling can give rise to a Sender/Receiver hybrid phenotype. Proc National Acad Sci. 2015;112: E402–E409. doi:10.1073/pnas.1416287112

52. Riahi R, Sun J, Wang S, Long M, Zhang DD, Wong PK. Notch1–Dll4 signalling and mechanical force regulate leader cell formation during collective cell migration. Nat Commun. 2015;6: 6556. doi:10.1038/ncomms7556

53. Alhashem Z, Feldner-Busztin D, Revell C, Portillo MA-G, Camargo-Sosa K, Richardson J, et al. Notch controls the cell cycle to define leader versus follower identities during collective cell migration. Elife. 2022;11. doi:10.7554/elife.73550

54. Stoker M, Gherardi E, Perryman M, Gray J. Scatter factor is a fibroblast-derived modulator of epithelial cell mobility. Nature. 1987;327: 239–242. doi:10.1038/327239a0

